# CeSpGRN: Inferring cell-specific gene regulatory networks from single cell multi-omics and spatial data

**DOI:** 10.1101/2022.03.03.482887

**Authors:** Ziqi Zhang, Jongseok Han, Le Song, Xiuwei Zhang

## Abstract

Single cell profiling techniques including multi-omics and spatial-omics technologies allow researchers to study cell-cell variation within a cell population. These variations extend to biological networks within cells, in particular, the gene regulatory networks (GRNs). GRNs rewire as the cells evolve, and different cells can have different governing GRNs. However, existing GRN inference methods usually infer a single GRN for a population of cells, without exploring the cell-cell variation in terms of their regulatory mechanisms. Recently, jointly profiled single cell transcriptomics and chromatin accessibility data have been used to infer GRNs. Although methods based on such multi-omics data were shown to improve over the accuracy of methods using only single cell RNA-seq (scRNA-seq) data, they do not take full advantage of the single cell resolution chromatin accessibility data.

We propose CeSpGRN (**Ce**ll **Sp**ecific **G**ene **R**egulatory **N**etwork inference), which infers cell-specific GRNs from scRNA-seq, single cell multi-omics, or single cell spatial-omics data. CeSpGRN uses a Gaussian weighted kernel that allows the GRN of a given cell to be learned from the sequencing profile of itself and its neighboring cells in the developmental process. The kernel is constructed from the similarity of gene expressions or spatial locations between cells. When the chromatin accessibility data is available, CeSpGRN constructs cell-specific prior networks which are used to further improve the inference accuracy.

We applied CeSpGRN to various types of real-world datasets and inferred various regulation changes that were shown to be important in cell development. We also quantitatively measured the performance of CeSpGRN on simulated datasets and compared with baseline methods. The results show that CeSpGRN has a superior performance in reconstructing the GRN for each cell, as well as in detecting the regulatory interactions that differ between cells. CeSpGRN is available at https://github.com/PeterZZQ/CeSpGRN.

## 1. Introduction

Gene regulatory networks (GRNs) represent how genes regulate each other during biological processes. Inferring GRNs from gene expression data has been a long-standing and challenging problem. Single-cell gene expression data have been used to infer GRNs, where each cell is used as a sample [1, 2, 3, 4, 5, 6, 7]. These methods aim to learn one GRN from the gene expression data of a population of single cells from either a cell cluster or all the cells in one dataset. However, it has been reported that GRNs are highly dynamic and their topologies change over a temporal process or a spatial landscape [8, 9, 10, 11, 12, 13]. Each cell can have its unique GRN according to its developmental stage along the dynamic process. Obtaining cell-specific GRNs is of great significance as it allows researchers to investigate the changes in GRNs during a dynamic process or cross a spatial landscape. Dictys [14] was recently proposed to infer time-varying GRNs along the differentiation trajectory using single-cell gene expression data. The method moves a window along the trajectory and infers the GRN for each window step. However, the method does not infer GRNs at single cell resolution, which fails to capture detailed rewiring processes along the trajectory. Furthermore, Dictys requires that cells are pre-ordered by pseudotime, but the pseudotime inference step can also introduce errors. Dai *et al* developed CSN (Cell Specific Network), which attempts to calculate pairwise gene-gene association in single cells using a statistical measurement [15]. However, CSN analyzes the association for each gene-pair independently. In GRN inference all genes are considered at the same time, which is more challenging than pairwise gene-gene association analysis.

Meanwhile, the advance of single-cell multi-omics technology makes it possible to utilize the cross-modalities information to infer better GRN. Methods have been proposed to use single-cell chromatin accessibility and gene expression data (e.g. scATAC-seq and scRNA-seq data) to infer GRN [16, 17, 18]. Chromatin accessibility information of transcription factor (TF) binding sites is used to provide a “prior graph” that helps to narrow down the regulatory events between TFs and target genes. However, these methods still focus on inferring population-level GRN and they only use bulk-level peak information in scATAC-seq data to construct the prior graph. Spatial transcriptomics data provides the spatial location of cells along with the gene expression data. Spatially adjacent cells tend to have stronger interactions compared to distant cells, and methods have been proposed to infer cell-cell interactions (CCIs) from spatial transcriptome data [19, 20, 21]. Inferring cell-specific GRNs in spatial data can conceptually lead to better prediction of CCIs by connecting CCIs and within-cell interactions (GRNs).

In this paper, we propose CeSpGRN (**Ce**ll **Sp**ecific **G**ene **R**egulatory **N**etwork inference), a computational method that infers cell-specific GRNs from single-cell multi-omics and spatial data. Typical GRN inference methods require a sufficient number of cells to infer one GRN, which makes inferring cell-specific GRNs for all cells seemingly impossible. In CeSpGRN, we assume that the GRNs of cells change smoothly along the cell trajectory or spatial landscape. This assumption enables “information borrowing” across different cells: when inferring the GRN for a cell, CeSpGRN uses data from not only the cell itself, but also its neighboring cells. The neighboring cells in CeSpGRN are defined according to the similarity of gene expression data or spatial location. With the “information borrowing” strategy, CeSpGRN does not require additional trajectory inference or cell clustering steps that was used in many high-resolution GRN inference methods [14, 22], and circumvent the potential error introduced by these steps. CeSpGRN uses a Gaussian Copula Graphical Model (GCGM) [23, 24] to model the gene expression data. GCGM is expanded upon the Gaussian Graphical Model (GGM). Compared to GGM, GCGM account for the non-Gaussian nature of single-cell gene expression data. The design of CeSpGRN makes it possible to apply the method to single-cell multi-omics datasets. Given the paired scATAC-seq and scRNA-seq data where both modalities are simultaneously measured for each cell, CeSpGRN learns the cell-specific prior GRNs from the scATAC-seq data and refine the prior GRNs with the scRNA-seq data. When scATAC-seq data is not available, CeSpGRN constructs prior GRNs from transcription factor information instead. Given the spatial transcriptomics data, CeSpGRN can also incorporate the spatial information into the weighted kernel construction.

We tested CeSpGRN on 3 real datasets covering difference GRN inference scenarios, including a paired scATAC-seq and scRNA-seq dataset, a spatial transcriptomics dataset, and a scRNA-seq dataset. We also quantitatively measure the performance of CeSpGRN on simulated datasets. The test result shows the broad applicability of CeSpGRN and its superior performance in reconstructing dynamically rewired regulations.

## 2. Methods

### 2.1. Overview of CeSpGRN

CeSpGRN can be separated into two steps: (1) construction of kernel weights, and (2) inference of cell-specific GRNs using kernel weights (Fig. 1). CeSpGRN is built upon the basic assumption that the cells’ GRNs change smoothly along the cell trajectory or spatial landscape, and the cells that are close to each other should have similar GRNs (termed “smooth-changing” assumption). CeSpGRN first constructs a *k*-nearest neighborhood (*k*-NN) graph. If the “smooth-changing” assumption is built upon the cell trajectory, the *k*-NN graph is constructed using gene expression data. If the assumption is built upon the spatial landscape (spatial transcriptomics data), then *k*-NN graph is constructed using spatial location of cells. The kernel weights are then calculated by applying a Gaussian kernel function on the *k*-NN graph (Sec. 2.2). The kernel weight between cells *i* and *j* is denoted as **K**_*ij*_.

**Figure 1.**
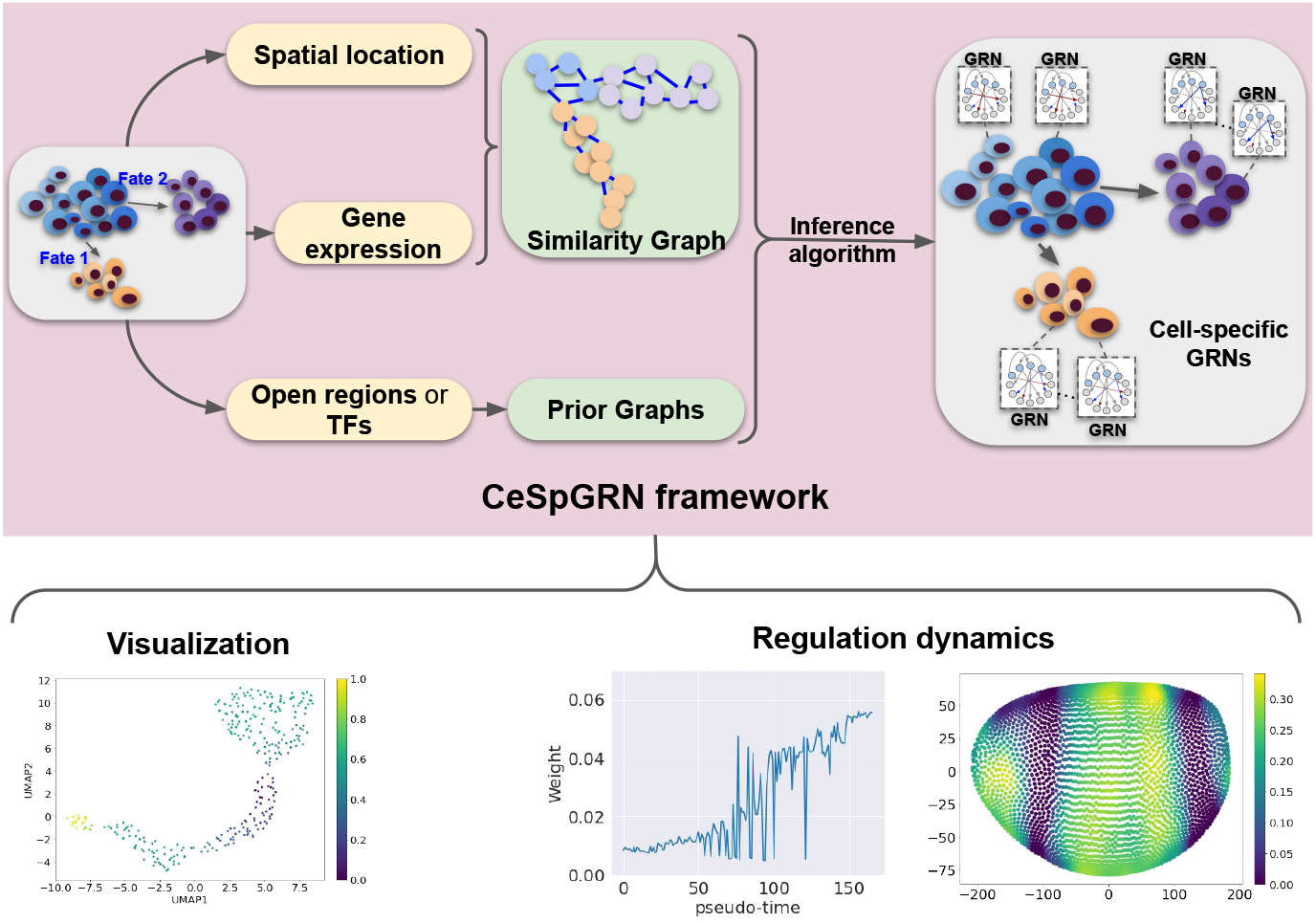
Overview of CeSpGRN framework.

We then construct the objective function of CeSpGRN using the kernel weights. For cell *i*, given the gene expression data **X**_*i*_ and the corresponding GRN **Θ**_*i*_ (belonging to the positive definite set 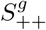), the likelihood function can be written as 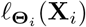. According to the “smooth-changing” assumption, the expression value of cell *j* from the neighborhood of cell *i* (*j* ∈ *N*_*i*_) should also partially reflect the GRN **Θ**_*i*_. We denote the likelihood of observing **X**_*j*_ given **Θ**_*i*_ as 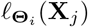. The objective function of cell *i* is a weighted average of the likelihood function of itself and its neighboring cells:

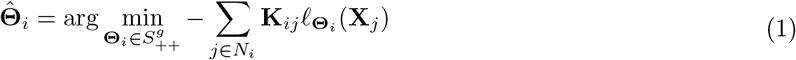

If cell *j* lies further to cell *i*, then the gene expression of cell *j* should reflect less of **Θ**_*i*_. This is controlled by a smaller kernel weight **K**_*ij*_ that is applied on 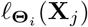 in the weighted average. Similarly, cell *j* lies closer to cell *i*, it should reflect more of **Θ**_*i*_, which means a larger kernel weight. The likelihood function 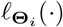 is calculated using Gaussian Copula Graphical Model (Sec. 2.4). When the chromatin accessibility data or TF information are available for the cells, cell-specific prior GRN can be constructed and incorporated into the objective function (Sec. 2.3). Denoting the binary adjacency matrix of prior GRN in cell *i* as 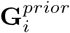, we can calculate the mask matrix of cell *i* by conducting an element-wise reverse: 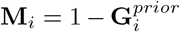. We further incorporate the *ℓ*_1_ regularization on **Θ**_***i***_ to control the sparsity of the inferred graph. The final objective function can be written as

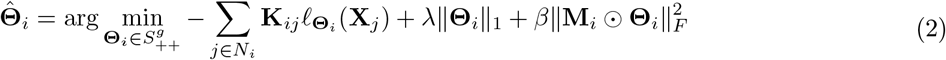

*λ*∥**Θ**_*i*_∥_1_ is the *ℓ*_1_ sparsity regularization on **Θ**_***i***_ and *λ* is the hyper-parameter. **M**_*i*_ ⊙ **Θ**_*i*_ denotes the element-wise multiplication of **M**_*i*_ and **Θ**_*i*_, and the term 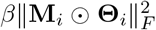 penalizes the difference between the prior and inferred GRNs. *β* is the regularization weight that controls the strength of this constraint.

The objective function is solved separately for each cell *i* in the dataset using ADMM algorithm (Sec. 2.4). The values in the inferred **Θ**_*i*_ cannot be readily used as the edge weights of the GRN. We further transform **Θ**_*i*_ into a partial correlation matrix **G**_*i*_ (Sec. 2.5), where a larger absolute partial correlation value at the (*m, n*)th element of **G**_*i*_ means a stronger regulation interaction between genes *m* and *n*.

The hyper-parameters in CeSpGRN mainly include the kernel bandwidth *σ*, neighborhood size *N*_*i*_, sparsity regularization weight *λ*, and prior information regularization weight *β*. We discuss the hyper-parameters selection in detail in Supplementary Note S2.3.

### 2.2. Constructing kernel weighted matrix K_*ij*_

To enforce the “smooth-changing” assumption on cell trajectory, the kernel weight **K**_*ij*_ should be constructed to reflect the transcriptome similarity between cell *i* and cell *j*. We adapt the idea from manifold learning [25, 26] to calculate **K**_*ij*_ for every pair of cells. We first perform Principal Component Analysis (PCA) on the scRNA-seq data matrix, and calculate pairwise Euclidean distance between cells using their low-dimensional PCA representations. We then construct a *k*-NN graph between cells using the pairwise distance matrix. We are then able to approximate the manifold distance between every two cells by calculating the corresponding geodesic distance [27] between them on the *k*-NN graph. Denote the geodesic distance between cells *i* and *j* by **D**_*ij*_, the kernel weight **K**_*ij*_ is then calculated using a Gaussian kernel function:

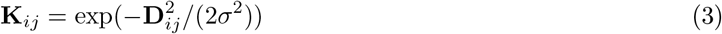

where *σ* is the bandwidth hyper-parameter that accounts for the differences between cell GRNs. A larger *σ* means that GRN is changing slower between cells, and a smaller *σ* means that GRN is changing faster. The number of neighbors in the *k*-NN graph, *k*, is set to be the minimum number larger than 5 that makes the graph connected. We select this *k* value because a *k* value that is too large makes the estimated manifold distance vulnerable to short-circuit errors, and a *k* value that is too small tends to fragment the data manifold into disconnect regions [28].

When dealing with spatial transcriptomics data, the “smooth-changing” assumption is applied on spatial landscape of cells. In this case, **D**_*ij*_ is the Euclidean distance between the spatial locations of cells, and the rest of the procedure to calculate **K**_*ij*_ remains the same.

### 2.3. Constructing 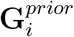 using chromatin accessibility data or TF knowledge

When the scATAC-seq data of each cell is provided, we can construct prior GRNs for each cell using the chromatin accessibility information. Given an accessible chromatin region, we can connect the region to the target gene using the relative location between regions and genes on the genome. In our test, we consider the regions that lie within 50kb upstream of the target gene TSS to be the regions that connect to the target gene. On the other hand, the region can be connected to the transcription factor using the motif information in the region. According to the accessible regions, we can connect the transcription factors to the corresponding target genes. We term the constructed graph between transcription factors and target genes to be the prior GRN. Different sets of regions are accessible in different cells. As a result, the constructed prior GRN is also cell-specific (Fig. S1). We construct the binary adjacency matrix of the prior GRN in cell *i* as 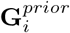, where 0 element means the interaction does not exist at the corresponding entry and 1 otherwise.

When the paired scATAC-seq data is not provided, CeSpGRN can still construct prior GRN using the TF information, *i*.*e*., which genes are TFs and which genes are target genes. When transforming the TF information into the prior GRN 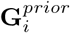, we set the entries corresponding to the interaction between target genes to be 0, and the remaining entries to be 1 (TF & TF, TF & Target). That is because the interactions in GRN are mainly between TF and TF, or TF and the target gene.

### 2.4. Inferring cell-specific GRNs using weighted GCGM

In this section, we introduce Guassian Copula Graphical Model (GCGM) and the detailed objective function of CeSpGRN (Eq. 2). GCGM expands upon Gaussian Graphical Model (GGM). GGM assumes that the data follows a multivariate Gaussian distribution, and the underlying graph is encoded in the precision matrix of the model. However, GGM is limited to data with Gaussian distribution. GCGM is then developed to better account for the non-Gaussian nature of the data. In GRN inference, we assume the gene expression data is observed from GCGM, and we aim to infer the precision matrix that corresponds to the GRN. Inferring the precision matrix of a GGM or GCGM from the observed data has been studied by previous work [29, 30, 31, 23, 24].

Denoting the gene expression data of cell *i* as a vector **X**_*i*_ ∈ ℝ^*g*^, where *g* is the number of genes, and the precision matrix of the undirected GRN as **Θ**. The log likelihood function of GCGM can be written as:

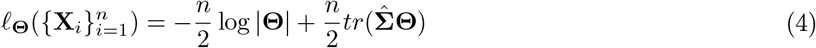

*n* is the number of cells, and 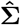 is the *non-parametric covariance matrix* of GCGM. We then include the log-likelihood function of GCGM (Eq. 4) into the objective function of CeSpGRN (Eq. 2):

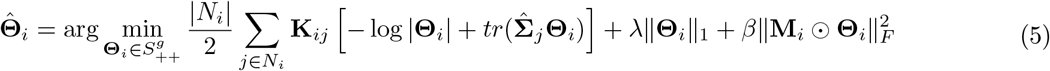

The remaining problem is the estimation of *nonparametric covariance matrix* 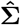. In the original GCGM model, 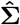 is calculated based on Kendall’s *τ* correlation [23, 24]. In CeSpGRN, we need to calculate 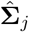for every cell *j*. We again borrow information from neighboring cells of cell *j*, and estimate 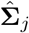 following:

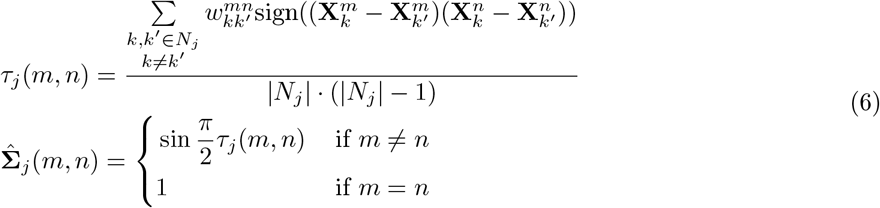

where 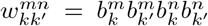, and 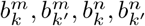 are binary annotation for the zero values. 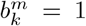 if the expression of gene *m* in cell 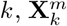, is not zero, and 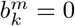 if 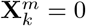. The estimated 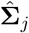 is not guaranteed to be positive definite, and we project 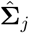 into the positive definite sets by replacing its non-positive eigenvalues with a small positive value *ϵ* = 10^−3^. *N*_*j*_ is the set of neighboring cells set of cell *j*. The size of neighborhood |*N*_*j*_| is a hyper-parameter in the formula, with a similar role as that of the bandwidth parameter *σ*.

Having calculated kernel weight **K**_*ij*_ and 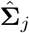, we now can proceed to solve the optimization problem (Eq. 5). We use the ADMM algorithm to perform the optimization, as (1) ADMM preserves the positive definiteness of **Θ**_*i*_ for each iteration as long as the initial **Θ**_*i*_ is positive definite (proof in Supplementary Note S2.2); (2) ADMM quickly converges to a reasonably good suboptimum [32]; (3) The convergence condition and hyper-parameter choices for the algorithm are well-studied [32, 33]. To apply ADMM, we transform the original optimization problem into the following problem:

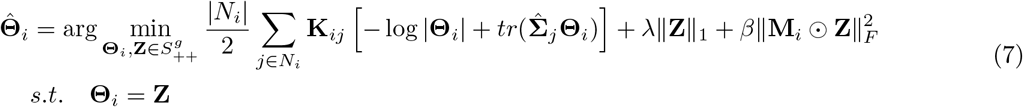

The new optimization problem in Eq. 7 can be solved with `Algorithm 1` (pseudo-code in Supplementary Note S2.1).

### 2.5. Calculating partial correlation matrix

The zero entries in the inferred precision matrix **Θ**_*i*_ means conditional independence of the corresponding genes. However, the non-zero values of **Θ**_*i*_ cannot be directly used to quantify the regulation strength of genes. We introduce an additional step to transform the precision matrix **Θ**_*i*_ into the partial correlation matrix **G**_*i*_. Given the (*m, n*)th entry of **Θ**_*i*_, the corresponding **G**_*i*_(*m, n*) can be calculated as

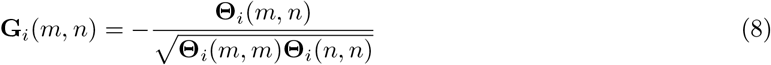

The values of the partial correlation matrix reveal the regulation strength between genes, where a larger value means the corresponding genes have a stronger regulation relationship.

## 3. Results

### 3.1. CeSpGRN infers cell-specific GRNs from paired scRNA-seq and scATAC-seq dataset

We test the performance of CeSpGRN on a paired scATAC-seq and scRNA-seq dataset of mouse neuromesodermal progenitors (NMPs) differentiation [34]. The dataset measures both chromatin accessibility and gene expression data for each individual cell within a sample NMPs population. NMPs are a type of bipotent stem cell that can further differentiate into elongating spinal cord cells and somitic mesoderm cells [34]. Here we selected 300 cells from the dataset that cover both differentiation trajectories of NMPs (Fig. 2a), and selected 610 genes that include both highly-variable genes and key transcription factors. We constructed cell-specific prior GRNs from the scATAC-seq data (Sec. 2.3), and then ran CeSpGRN using the prior GRNs and gene expression data. We visualized the inferred GRN matrices using UMAP, and color each matrix using the cell type annotation of its corresponding cells in the original literature [34] (Fig. 2b). From the UMAP visualization, we observed that the inferred cell-specific GRN still preserves the bifurcating trajectory of NMPs, where one branch leads to the spinal cord cells (red cells in Fig. 2b) and the other branch leads to the somite mesoderm cell (green cells in Fig. 2b). We then analyzed the highly variable hub genes under each differentiation branch. The highly variable hub genes are selected according to the variance of the total edge weights that connect to each gene along the branch, and a larger variance means that the corresponding gene shows more significant rewiring activity in the differentiation process. We selected the top 100 hub genes for each branch and ran gene ontology analysis using topGO [35]. The top gene ontology terms of each branch are shown in Table. S1. Both branches include the term “anterior/posterior axis specification”, which is highly relevant to NMPs differentiation [36]. In the somite mesoderm branch, we also found a highly relevant GO term “mesoderm development”.

**Figure 2.**
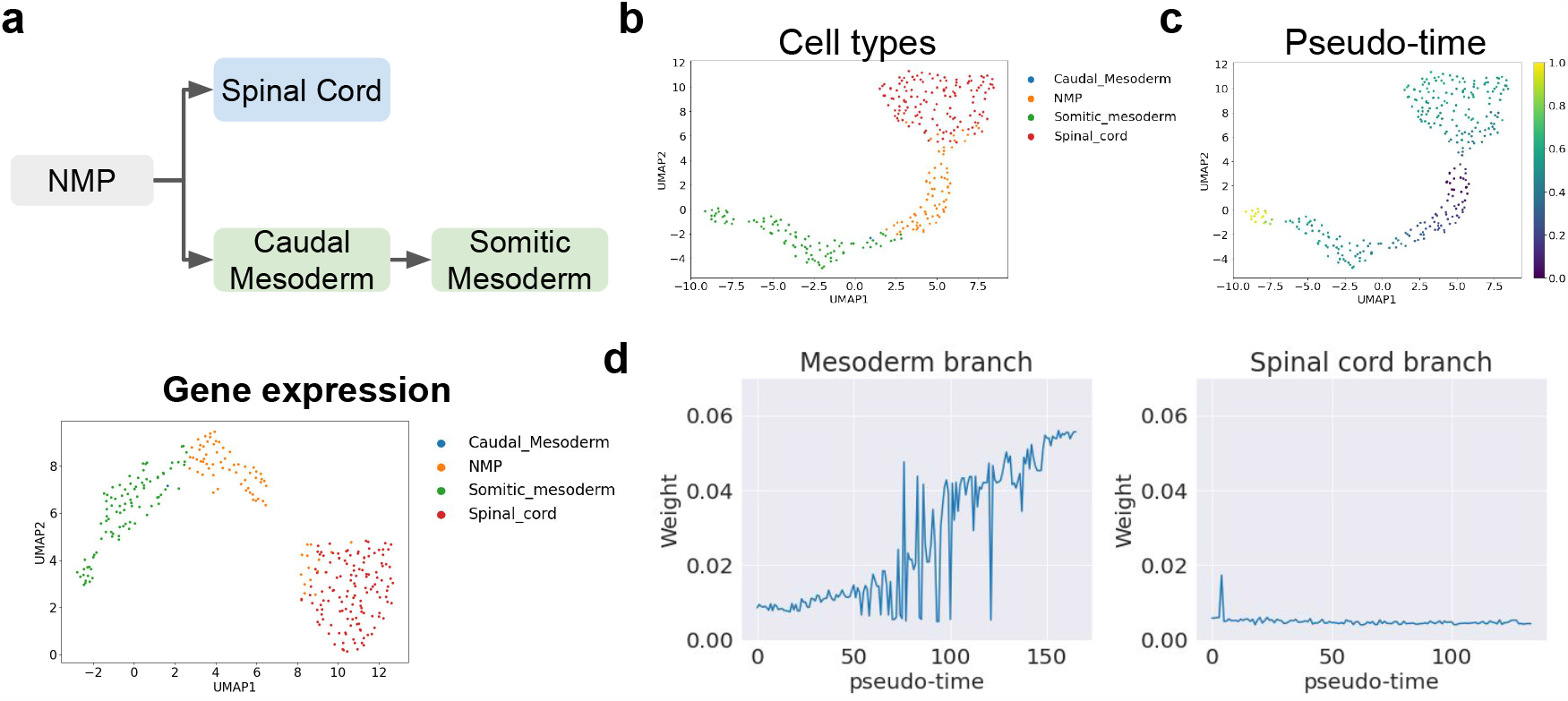
Test result on NMP differentiation dataset. **a**. (upper) The differentiation trajectory of NMP cells. (lower) the UMAP visualization of gene expression data. **b-c**. The UMAP visualization of inferred GRNs, where cells are colored by (b) the original cell type annotation, and (c) the diffusion pseudotime. **d**. The change of edge weight between *Mesp2* and *Ripply2* along the Mesoderm branch and Spinal cord branch.

Cells in mesoderm trajectory are affected by the regulation between genes *Mesp2* and *Ripply2*. During the somitogenesis process, the expression of *Mesp2* induces the expression of *Ripply2* which in return regulates the total abundance of *Mesp2*. The regulation results in the spatial periodicity of the segmented somites [34]. Here we analyzed the change in regulation weight between *Mesp2* and *Ripply2* along both branches. We first inferred the developmental pseudotime of each cell using diffusion pseudotime algorithm [37] (Fig. 2c). Then we sort the cells according to the inferred pseudotime along each branch, and plot the edge weight between *Mesp2* and *Ripply2* following the sorted order of cells (Fig. 2d). In the mesoderm branch, it can be clearly observed that the weight increase and becomes more significant in somitic mesoderm cells, whereas the weight remains to be 0 along the whole spinal cord branch. The result shows the branch egulation between *Mesp2* and *Ripply2*, which further validates the inferred GRNs.

### 3.2. CeSpGRN detect spatially-associated regulation changes from spatial transcriptomics data

CeSpGRN can also be applied to spatial transcriptomics data. By utilizing the spatial location of the cells, CeSpGRN is able to learn how the regulation dynamics between genes change over space. We applied CeSpGRN to a Drosophila embryo dataset [38]. The dataset measures the spatial transcriptome of 84 genes from 3039 cells. Instead of using gene expression data, we use the 3D location of cells in the space to construct the weighted kernel function in CeSpGRN (Sec. 2.2). We then infer the cell-specific GRN among the 84 genes with the gene expression measurement. Using the inferred GRN, we then study the spatial-specificity of gene regulation that governs the Drosophila embryo morphogenesis process. We first explore the key regulators that drive the developmental process. We calculate the total absolute edge weights that connect to each gene for each cell. We then average the total absolute edge weights across cells and rank the genes according to the average weights. Among the top-scoring genes, we found multiple transcription factors that were reported to drive the morphogenesis process, including multiple pair-rule genes *eve, odd, prd*, and *ftz* (Fig. 3a). We then selected the top 20 genes and conducted the gene ontology analysis using *topGO*. We discovered multiple terms related to embryo development and segmentation: “anatomical structure formation involved in morphogenesis”, “trunk segmentation”, etc (Fig. 3b). We then explore the edges that rewire the most across space. We calculate the variance of edge weights for each edge in the inferred GRNs, and rank the edges according to their variances. We found highly variable edges between *ftz* and *eve, eve* and *zen, odd* and *eve*, etc. Interestingly, most of the highly-variable edges includes genes *eve* and *ftz*, and these two genes were reported to bind to most genes in Drosophila embryos in order to regulate the morphogenesis process [39] (Fig. 3c). We further select several highly-variable edges connecting to key regulators *eve, odd*, and *ftz*, and visualize the weight distributions of these edges across space (Fig. 3d). The segmentation of Drosophila embryo results in a more abrupt structural change over space. To capture a more detailed shift of regulation weight corresponding to the structural change, we used the edge weights from the GRNs inferred with a small bandwidth (0.01) instead of the aggregated GRNs (Supplementary Note S2.3), since aggregated GRNs include results from larger bandwidth that potentially over-smooth the regulation change. We visualize the spatial distribution of weights corresponding to edges “ftz-eve”, “ftz-odd”, and “eve-zen” in Fig. 3d. From the visualization, we observe a strong spatial specificity of those edges across space. *ftz* and *eve* are important transcription factors related to embryonic segmentation, and the distribution of their expression level also follows the anatomical segmentation structure of Drosophila embryo [39, 38] (Fig. 3e). We also observe a similar segmentation pattern in the weight distribution between *ftz* and *eve* (Fig. 3d left), which reveals that the regulation strength between “ftz-eve” could also follow the segmentation structure of the Drosophila embryo. Gene *zen* controls the development of dorsal-ventral pattern in Drosophila embryo, and it is mainly expressed in the dorsal part of the embryo [40] (Fig. 3d right). From the weighted distribution of “eve-zen” (Fig. 3d right), we also observed a similar pattern, where the regulation weights are stronger in the ventral part and weaker in the dorsal part. With the inferred GRNs, we observed that the spatial specificity of expression for key regulators usually connects to the shift of the corresponding regulation weights. More experiments can be designed to further validate such correspondence.

**Figure 3.**
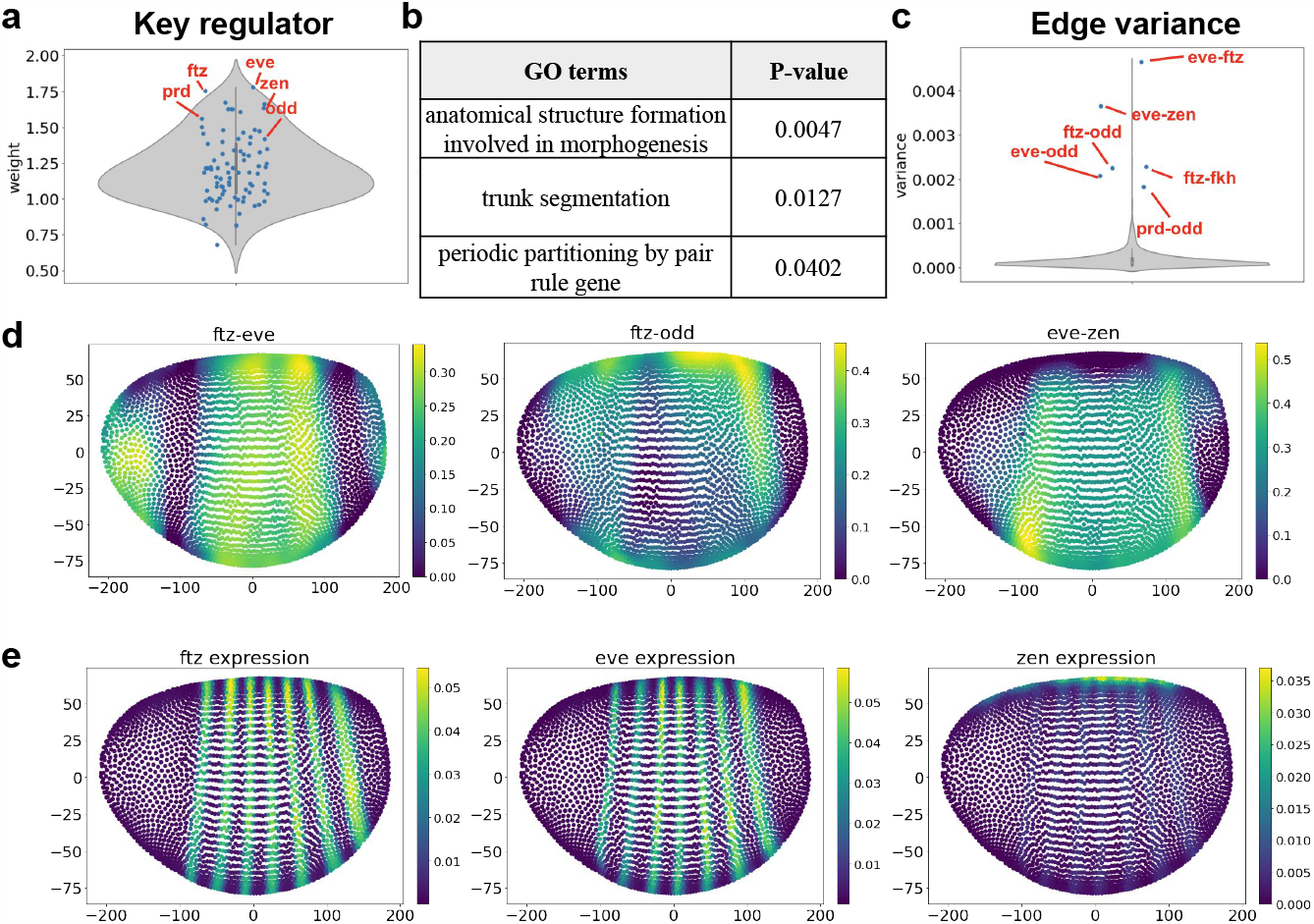
Test result on Drosophila embryo dataset. **a**. Violin plot showing the total weights for each genes in the dataset. **b**. The gene ontology term corresponds to the key regulators with the largest weight. **c**. Violin plot showing the distribution of edge variance across the space. **d**. The spatial distribution of edge weights, including edges “ftz-eve”, “ftz-odd”, and “eve-zen”. **e**. The spatial distribution of gene expression, including genes *ftz, eve*, and *zen*.

### 3.3. CeSpGRN detects regulation dynamics in mouse embryonic stem cells

We further applied CeSpGRN to a mouse embryonic stem cells (mESC) dataset [41]. In this dataset, 2717 mESC were sequenced using scRNA-Seq technology. The dataset includes cells from the following developmental stages: “before leukemia inhibitory factor (LIF) withdrawal”, “2 days after LIF withdrawal”, “4 days after LIF withdrawal”, and “7 day after LIF withdrawal”. These four stages correspond to four cell types respectively labeled as “mES cells”, “day 2”, “day 4” and “day 7”. With the LIF withdrawal, cells start to differentiate. The dataset records the onset of differentiation where cells fluctuate between a pluripotent inner cell mass-like (ICM) state to a differentiating epiblast-like state.

We preprocessed the dataset and selected 96 key genes which are highly variable genes and are also included in the key regulatory gene list reported in [42]. Among the 96 genes, 10 are known to be TFs (*Pou5f1, Nr5a2, Sox2, Sall4, Otx2, Esrrb, Stat3, Tcf7, Nanog, Etv5*) and the remaining are target genes [42]. The UMAP visualization of the preprocessed gene expression data is shown in Fig. 4a. We constructed prior GRNs using the TF information and inferred cell-specific GRNs using CeSpGRN (Sec. 2.3). We visualized the inferred GRNs using UMAP (Fig. 4b). From the UMAP visualization, we observe a clear differentiation trajectory matching the trajectory pattern of gene expression data (Figs. 4a,b).

**Figure 4.**
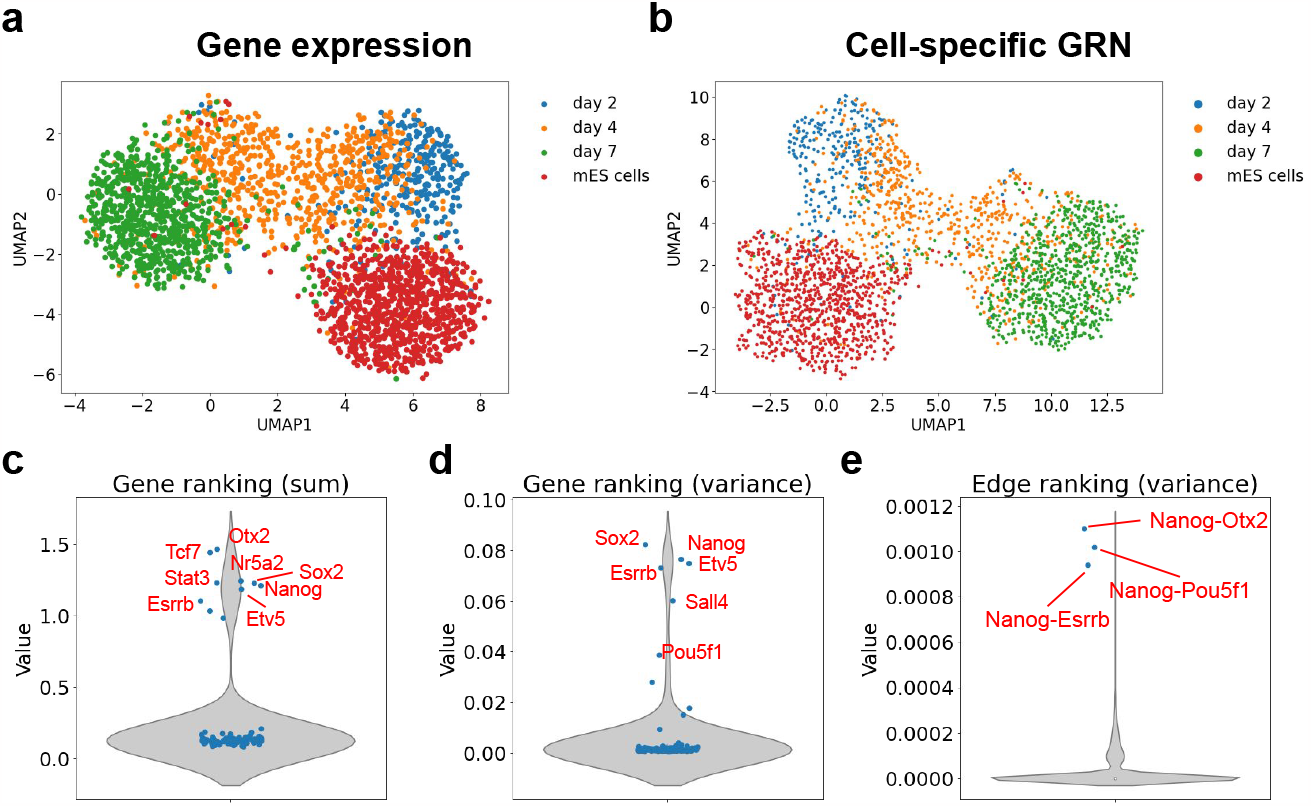
(a) UMAP visualization of gene expression data. Cells are colored by the cluster they are associated with. (b) UMAP visualization of cell-specific GRNs inferred by CeSpGRN-TF. Dots are colored according to the cells’ labels. Legend is the same as the one used in (a). (c) Violin plot showing the total edge weight connecting to each gene. (d) Violin plot showing the variance of edge weight connecting to each gene. (e) Violin plot showing the variance of edge weights.

The cell-specific GRNs inferred by CeSpGRN allow us to analyze the dynamics of GRNs in this dataset. First, we calculate the means and variances of total edge weights connecting to each gene across all inferred GRNs. We sort the genes according to their mean and variance values separately. The mean and variance measure two different aspects of genes: a larger mean total edge weight means that the gene potentially has more regulation activities, whereas a larger variance means that the gene rewires more often in the cell population. We observed several genes that rank at the top in both evaluating metrics, including *Otx2, Tcf7, Nanog*, etc. These genes were shown to play crucial roles in mESC development [43, 42] (Figs. 4c,d). We also check the variance of edge weights and select the edges that vary the most in the cell population. CeSpGRN detected top-ranking edges including *Pou5f1* (*Oct4*)-*Nanog, Esrrb*-*Nanog*, and *Nanog* -*Otx2* (Fig. 4e). Those edges were discussed as the key interactions for mESC differentiation literature [42, 44, 45, 46].

We further check how the total number of target genes of the key regulators change along the differentiation trajectory. We infer the cell developmental pseudotime using diffusion pseudotime (DPT) algorithm [37]. For each cell along the trajectory, we then calculate the total number of target genes connecting to the key regulators by thresholding the edge weights. We then plot the target gene number of regulators *Nanog* and *Sox2* along the trajectory (Figs. 5a,b). For both genes, we observed smooth changes in edge weights as cells fluctuate between ICM-like state and epiblast-like state. Interestingly, we observe that for both genes, the initial target gene numbers are much smaller in “mES cells” cluster and increase gradually along the differentiation path. The trend also matches the experiments, where LIF is withdrawn after “mES cells” and cells start to differentiate.

**Figure 5:**
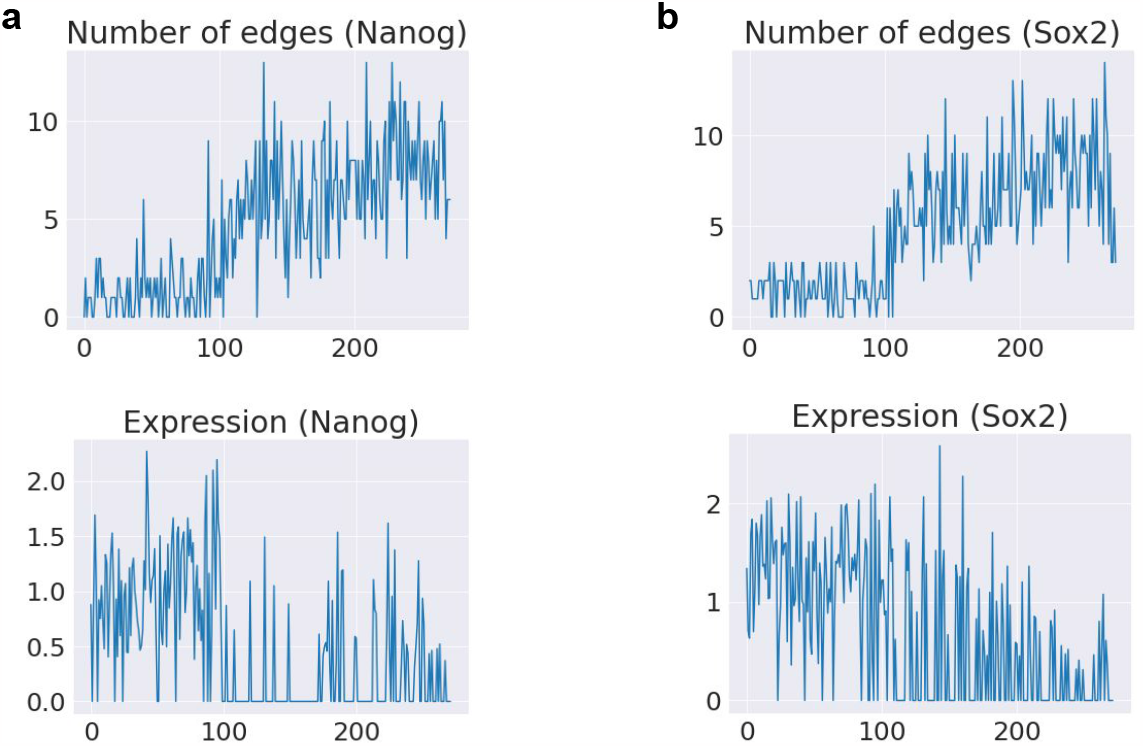
(a) The change of (upper) total edge number connecting to *Nanog* and (lower) *Nanog* expression level along the trajectory. (b) The change of (upper) total edge number connecting to *Sox2* and (lower) *Sox2* expression level along the trajectory.

### 3.4. Benchmark on simulated datasets

We further benchmark the inference accuracy of CeSpGRN on simulated datasets, and compare it with baseline GRN inference methods including scMTNI [22], GENIE3 [47], and CellOracle [48]. scMTNI is a recently proposed method that infers cluster-level GRNs using scATAC-seq and scRNA-seq data. When scATAC-seq is not available, scMTNI infers GRNs using scRNA-seq data and TF information. Here we denote the scMTNI that uses TF information and scATAC-seq data as “scMTNI (TF)” and “scMTNI (ATAC)”, respectively. CellOracle is a regression model that infers cluster-level GRNs using scATAC-seq and scRNA-seq data. GENIE3 is a tree-based inference method that infers population-level GRNs from scRNA-seq data and TF information (termed “GENIE3 (TF)”). In a recent benchmarking paper [1], GENIE3 is among the top-performing methods that infer GRN from scRNA-seq data. We have selected these state-of-the-art cluster-level or population-level GRN inference methods as the baseline methods for comparison. The details of running the baseline methods are described in Supplementary Note S2.4. Next, we generate simulated datasets using scMultiSim [49], a simulation method that can simulate scRNA-seq data and scATAC-seq data from dynamically changing GRNs. The simulation details is included in Supplementary Note S2.5.

We first tested the methods under the scenario where only scRNA-seq data and the TF information are available. We generated 3 scRNA-seq datasets with different random seeds (0, 1, 2), and then benchmarked the performance of CeSpGRN (denoted as “CeSpGRN (TF)”) and two baseline methods including scMTNI (TF) and GENIE3 (TF). We measured the inference accuracy using AUPRC score and Early Precision scores (Supplementary Note S2.5), and plot the result in Fig. 6a. In the result, we observed that CeSpGRN (TF) has higher AUPRC and Early Precision scores, which shows that CeSpGRN (TF) better recovers the dynamically changed GRNs from scRNA-seq data.

**Figure 6.**
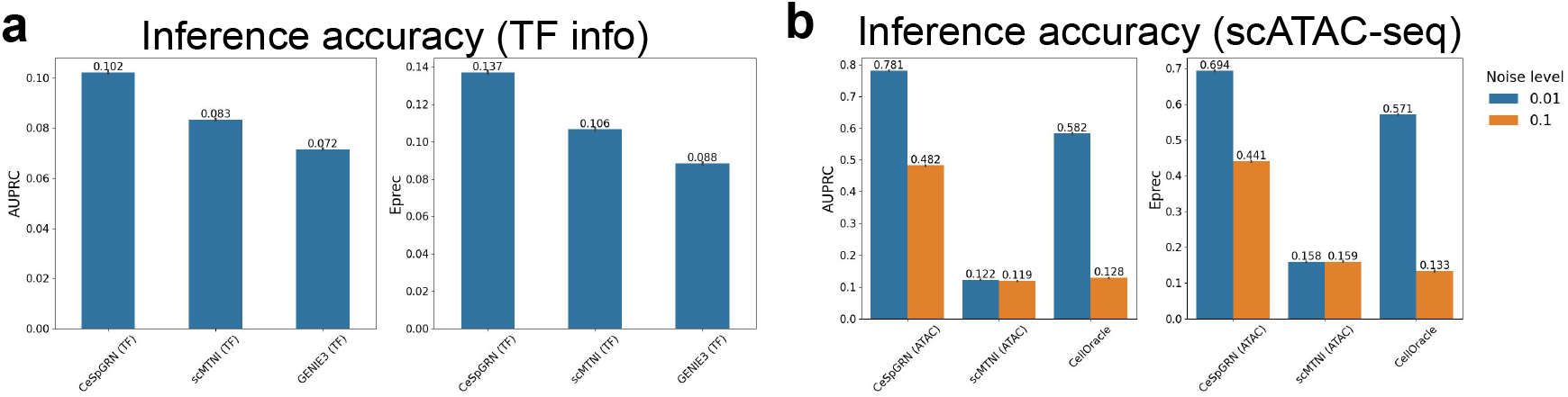
Test results on simulated datasets. **a**. The AUPRC and early precision (Eprec) scores of CeSpGRN and baseline methods when TF information is provided. **b**. The AUPRC and early precision (Eprec) scores of CeSpGRN and baseline methods when scATAC-seq is provided.

We further tested the methods under the scenario where both scRNA-seq data and scATAC-seq data are available for each cell. We generated 6 paired scRNA-seq and scATAC-seq datasets with different random seeds (0, 1, 2) and noise levels of cross-modalities relationship (0.01 and 0.1, Supplementary Note S2.5). The result (Fig. 6b) shows that CeSpGRN (ATAC) performs significantly better compared to baseline methods. This is because CeSpGRN (ATAC) better utilizes the accessible regions in each cell for cell-specific prior network construction, whereas the other methods only uses the peak information in scATAC-seq data. On the other hand, the noise levels of cross-modalities relationship also affect the overall performance of CeSpGRN (ATAC) (Fig. 6b). This is because a higher noise level means that the prior networks constructed from scATAC-seq data is less accurate, which finally affect the overall performance of CeSpGRN.

The benchmark result on simulated datasets jointly show that CeSpGRN is able to better capture the dynamical changes of GRNs, especially when scATAC-seq data are provided.

## 4. Discussion and future work

We proposed CeSpGRN, a method that can infer cell-specific GRNs from single-cell multi-omics or spatial data. The assumption that closely located cells (gene expression space or spatial landscape) have similar GRNs can significantly reduce the parameter space of the cell-specific GRNs inference problem. CeSpGRN constructs objective functions for each cell using a weighted Gaussian Copula Graphical Model, where the weight is calculated from either the gene expression value or the spatial location of the cells. When the paired scATAC-seq data is available, CeSpGRN innovatively constructs the cell-specific prior GRNs, and uses the prior GRNs to improve the accuracy of GRNs. Traditional GRN inference methods construct prior GRN only using the bulk-level peak information, which ignores the region accessibility of each cell in scATAC-seq data. The construction of cell-specific prior GRNs, on the other hand, fully explores scATAC-seq data by using both the peak information and the region accessibility of cells. The design of CeSpGRN allows it to be broadly applied to scRNA-seq data, paired scRNA-seq and scATAC-seq data, and spatial transcriptomics data. We demonstrate CeSpGRN using real and simulated datasets that cover various application scenarios. The test results show that CeSpGRN has a superior performance in detecting key regulators and regulations that rewired the most in cell distribution, and learning the trend of regulation changing, etc. CeSpGRN, being the first method that infers cell-specific GRNs using single-cell multi-omics data, also has its limitations. CeSpGRN infers undirected graphs whereas the regulations are directional in reality. The hyper-parameter still needs tuning sometimes, which a common issue that lies within most of the unsupervised GRN inference methods. However, with CeSpGRN serving as the starting point, we envision that more methods will be proposed to deal with these limitations.

## 5. Acknowledgements

This work was supported in part by the US National Science Foundation DBI-2019771 and National Institutes of Health grant R35GM143070.

## Supplementary Materials

### S1. Supplementary Figures and Tables

**Figure S1:**
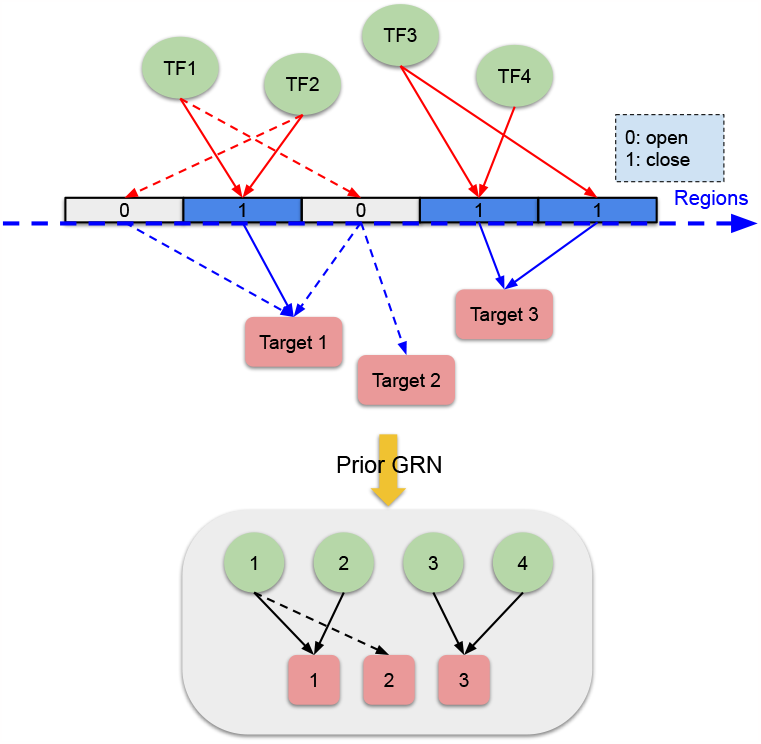
Graphical illustration of constructing prior GRN from chromatin accessibility data within a cell. Connections between TFs and targets are only preserved when the connecting regions are accessible. Solid lines: connections are preserved. Dashed lines: connections are removed since the regions are closed.

**Table S1:** The top gene ontology terms and the corresponding p-values of two branches (Spinal Cord branch and Mesoderm branch).

| GO terms | P-value | Branch |
| --- | --- | --- |
| anterior/posterior axis specification | 6.5e-5 | Mesoderm |
| embryonic axis specification | 0.0269 | Mesoderm |
| mesoderm development | 0.0488 | Mesoderm |
| anterior/posterior axis specification | 0.011 | Spinal Cord |
| embryonic axis specification | 0.024 | Spinal Cord |

### S2. Supplementary Note

### S2.1. Pseudo-code of CeSpGRN

#### Algorithm 1

CeSpGRN algorithm

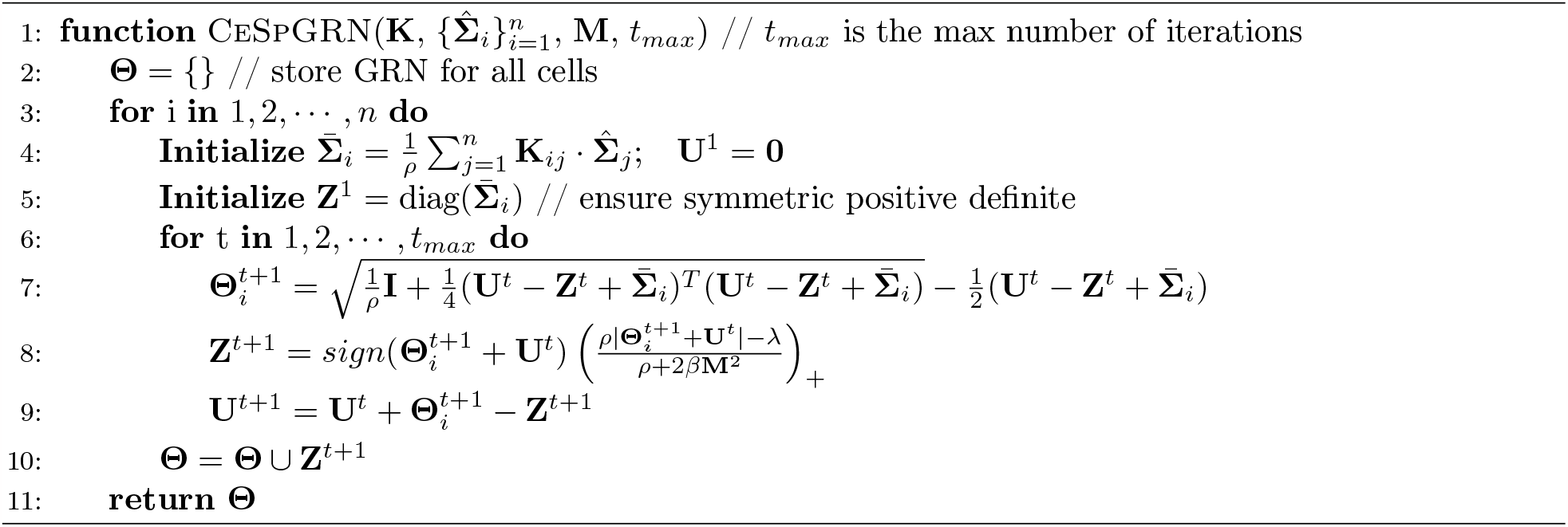

Parameter *ρ* in the algorithm affects the convergence speed of the algorithm, and is set to be 1.7 [33].

### S2.2. Algorithms 1 preserve the positive definiteness of the inferred matrix

When we initialize **Z**^0^ to be symmetric positive definite, then according to algorithm 1, we will always have **Z**^*t*^ to be symmetric. Then ^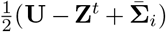^ is also symmetric, and can be eigenvalue decomposed as **VΛV**^*T*^, where **V**^*T*^ **V** = **VV**^*T*^ = **I**. Rewriting the updating rule of 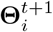 in Algorithm 1 as

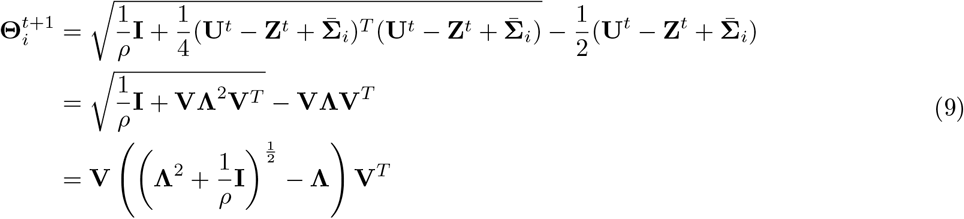

Since 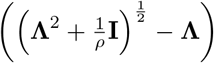 has minimum eigenvalue greater than 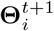 is always positive definite. Algorithm 1 is able to preserve the symmetric positive definiteness of 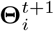.

### S2.3. Hyper-parameter selection

The hyper-parameters in CeSpGRN include the kernel bandwidth *σ*, neighborhood size *N*_*i*_, sparsity regularization weight *λ*, and prior information regularization weight *β*. The kernel bandwidth *σ* and neighborhood size *N*_*i*_ control the change speed of cell GRNs, where a smaller *σ* and *N*_*i*_ mean GRNs change faster across the cell population. *λ* controls how sparse the inferred GRN is, whereas *β* controls how close the inferred GRNs are to the prior GRNs. In our test, we fix *β* = 1 and *N*_*i*_ = 100. We scan through *σ* = [0.01, 0.1, 1, 10] and *λ* = [0.01, 0.05, 0.1, 0.5] and infer GRNs under each parameter setting. The final GRN is obtained by averaging the inferred GRNs under each parameter setting.

### S2.4. Running details of baseline methods

When running scMTNI, we first separate the cell population into 3 distinct clusters according to the simulation pseudotime. Then we ran scMTNI following the same pipeline in its online tutorial (https://github.com/Roy-lab/scMTNI). We set the argument “-q” to 5 when running the method with scATAC-seq data, which increase the influence of prior network on the final result. We adjust the “branch-specific gain/loss rate” in the cell lineage tree to 0.001 according to the ground truth GRN changing rate in the simulated datasets. When running CellOracle, we follow the same pipeline in its online tutorial (https://morris-lab.github.io/CellOracle.documentation/index.html). Same as scMTNI, we also separate the cell population into 3 clusters according to the simulation pseudotime.

### S2.5. Simulation setting and evaluation metrics

We generated simulated datasets using scMultiSim. Totally 3 scRNA-seq datasets (3 random seeds: 0, 1, 2) and 6 paired scRNA-seq and scATAC-seq datasets (3 random seeds: 0, 1, 2, and 2 noise levels: 0.01, 0.1) are generated. Each dataset has a total of 8000 cells and 110 genes. For paired scRNA-seq and scATAC-seq data, the noise level measures the amount of false positive connections in the cross-modality relationship matrices (region by TF matrix, region by target gene matrix). A higher noise level means that the relationship matrix is less accurate, which also affect the accuracy of prior GRNs since they are constructed from the cross-modality relationship matrix.

We evaluate the GRN inference accuracy of different methods using AUPRC score and Early Precision score. Both metrics were used in the previous benchmarking paper [1] to measure the closeness of the inferred GRNs towards the ground truth GRNs, where a higher score means that the inferred result is more accurate. Since CeSpGRN infers cell-specific GRNs, we measure the inference accuracy of each cell separately and average the scores across all cells. We also conducted the same cell-specific evaluation for the baseline methods to make the benchmarking result comparable across methods.

